# Mutation rate of SARS-CoV-2 and emergence of mutators during experimental evolution

**DOI:** 10.1101/2021.05.19.444774

**Authors:** Massimo Amicone, Vítor Borges, Maria João Alves, Joana Isidro, Líbia Zé-Zé, Sílvia Duarte, Luís Vieira, Raquel Guiomar, João Paulo Gomes, Isabel Gordo

**Affiliations:** Instituto Gulbenkian de Ciência, Oeiras, Portugal; Bioinformatics Unit, Department of Infectious Diseases, National Institute of Health Doutor Ricardo Jorge (INSA), Lisbon, Portugal; Centre for Vectors and Infectious Diseases Research, Department of Infectious Diseases, National Institute of Health Doutor Ricardo Jorge (INSA), Águas de Moura, Portugal; BioISI - Biosystems & Integrative Sciences Institute, Faculty of Sciences, University of Lisbon, Portugal; Innovation and Technology Unit, Department of Human Genetics, National Institute of Health Doutor Ricardo Jorge (INSA), Lisbon, Portugal; Centre for Toxicogenomics and Human Health (ToxOmics), Genetics, Oncology and Human Toxicology, Nova Medical School|Faculdade de Ciências Médicas, Universidade Nova de Lisboa, Lisbon, Portugal; National Reference Laboratory for Influenza and other Respiratory Viruses, Department of Infectious Diseases, National Institute of Health Doutor Ricardo Jorge (INSA), Lisbon, Portugal

## Abstract

**Background and objectives:** To understand how organisms evolve, it is fundamental to study how mutations emerge and establish. Here, we estimated the rate of mutation accumulation of SARS-CoV-2 *in vitro* and investigated the repeatability of its evolution when facing a new cell type but no immune or drug pressures.

**Methodology:** We performed experimental evolution with two strains of SARS-CoV-2, one carrying the originally described spike protein (CoV-2-D) and another carrying the D614G mutation that has spread worldwide (CoV-2-G). After 15 passages in Vero cells and whole genome sequencing, we characterized the spectrum and rate of the emerging mutations and looked for evidences of selection across the genomes of both strains.

**Results:** From the mutations accumulated, and excluding the genes with signals of selection, we estimate a spontaneous mutation rate of 1.25×10^-6^ nt^-1^ per infection cycle for both lineages of SARS-CoV-2. We further show that mutation accumulation is heterogeneous along the genome, with the spike gene accumulating mutations at rate five-fold higher than the genomic average. We also observe the emergence of mutators in the CoV-2-G background, likely linked to mutations in the RNA-dependent RNA polymerase and/or in the error-correcting exonuclease protein.

**Conclusions and implications:** These results provide valuable information on how spontaneous mutations emerge in SARS-CoV-2 and on how selection can shape its genome towards adaptation to new environments.

**Lay summary:** Mutation is the ultimate source of variation. We estimated how the SARS-COV-2 virus—cause of the COVID-19 pandemic—mutates. Upon infecting cells, its genome can change at a rate of 0.04 per replication. We also find that this rate can change and that its spike protein can adapt, even within few replications.

## Background and objectives

Mutation is the principal process driving the origin of genetic diversity. The mutation rate is a function of replication fidelity and represents the intrinsic rate at which genetic changes emerge prior to selection. The substitution rate, instead, is a measure of mutation accumulation in a given period of time and embeds the effects of selection[1]. These rates and the spectrum of the emerging mutations are fundamental parameters to understand how an organism evolves and how new variants are purged from, or establish in natural populations.

In DNA based microbes the genomic mutation rate per cell per generation, measured in laboratory conditions, is close to a constant[2]. On the other hand, for RNA viruses there is a remarkable variation in their replication fidelity[3,4]. The basic mutation rates, expressed as nucleotide substitutions per site per cell infection (s/n/c), vary between 10^-6^ to 10^-3^ for the several positive ssRNA viruses which have been studied[5]. Importantly, our current knowledge of the mutation rate of the human beta-coronavirus SARS-CoV-2, which is the cause of the COVID-19 pandemic[6], is based on estimates from different coronaviruses[5,7,8] and still lacks a direct quantification[9].

Laboratory evolution experiments with microbial populations allow to determine how fast mutations accumulate[10,11], and combining them with high-throughput sequencing is one of the best methods to estimate mutation rates, determine how they vary along the genome[12] and study the extent to which convergent evolution occurs[13,14].

Here, via experimental evolution of two natural variants of SARS-CoV-2[15] and whole genome sequencing, we characterized the spectrum and rates of their emerging mutations, and identified specific targets of selection. Such information is important for better understanding the basic biology of this virus and to quantify how predicable the evolution of strains with different transmission capabilities can be. It may also help determining some potential genomic constraints of the virus, which are key to the design of evolution proof vaccines and antiviral drugs.

## Methodology

### Virus growth and *in vitro* assay

Vero E6 (African green monkey, *Cercopithecus aethiops* kidney epithelial cells, ATCC^®^ CRL 1586™) cells were cultured at 37°C and 5% CO_2_ in Minimum Essential Medium (MEM 1X, Gibco^®^) supplemented with 10% fetal bovine serum (FBS), penicillin (100 units/ml) and streptomycin (100 μg/ml) + fungizone. The two clinical isolates Portugal/PT0054/2020 and Portugal/PT1136/2020, isolated at the National Institute of Health Doutor Ricardo Jorge (INSA), were used to produce the ancestors of the experimental evolution, CoV-2-D and CoV-2-G, which seeded the two laboratory evolution experiments. For this, the initial SARS-CoV-2 stock was produced by infecting Vero E6 cells (freshly grown for 24 h) and incubating the cells for 72 h. The culture supernatant was stored in aliquots at −80°C. The TCID_50_ of viral stock was calculated according to the method of Reed and Muench[16]. All work with infectious SARS-CoV-2 strains was done inside a class III microbiological safety cabinet in a containment level 3 facility at the Centre for Vectors and Infectious Diseases Research (INSA).

From the stored stocks, two 96-well plates fully inoculated with 50 μl of Vero E6 cells (2.0×10^4^ cells) grown for 24 h were infected with 50 μl of the SARS-CoV-2 strains viral suspension (2.0×10^3^ viruses) at a multiplicity of infection (MOI) of 0.1. MEM supplemented with 10% FBS, penicillin (100 units/ml) and streptomycin (100 μg/ml) + fungizone was added to each well (50 μl) and the plates were incubated for 24 h. Each well had a final volume of 150 μl. Every day, for 15 days, serial passages were done by passaging 50 μl of the culture supernatant to 96-well plates (one for each SARS-CoV-2 strain under study) fully inoculated with 50 μl of Vero E6 cells (2.0×10^4^ cells) using the same procedure and incubated in the same conditions. At day 15, total nucleic acids were extracted from 100 μl of viral suspension of each well in each plate (96 samples of day 15^th^ for each strain) using the automated platform NUCLISENS easyMAG (Biomérieux). Confirmation of nucleic acid integrity and rough concentration estimative was made before sequencing experiment by RT-qPCR of 8 random chosen samples from each plate at day 15 (CoV-2-D and CoV-2-G) using Novel Coronavirus (2019-nCoV) RT-PCR Detection Kit (Fosun Diagnostics). Samples from inoculation suspension (day 1) were also analyzed. All samples presented values of 7-10 Ct (Cycle threshold). When we infect the cells with 2×10^3^ particle forming units (PFU), after 24 h the number of PFUs is around 2×10^6^. So, assuming no major fluctuations in the viral load of the transferred suspension throughout the 15 passages and assuming a yield of approximately 1000 PFU/cell[9], the estimated number of replication cycles per passage is around 1 (i.e. ln(2×10^6^/2×10^3^)/ ln (10^3^)).

### SARS-CoV-2 genome sequencing and bioinformatics analysis

Genome sequencing was performed at INSA following an amplicon-based whole-genome amplification strategy using tiled, multiplexed primers[17], according to the ARTIC network protocol (https://artic.network/ncov-2019; https://www.protocols.io/view/ncov-2019-sequencing-protocol-bbmuik6w) with slight modifications, as previously described[15]. Briefly, after cDNA synthesis, whole-genome amplification was performed using two separate pools of tiling primers [pools 1 and 2; primers version V3 (218 primers) was used for all samples: https://github.com/artic-network/artic-ncov2019/tree/master/primer_schemes/nCoV-2019]. The two pools of multiplexed amplicons were then pooled for each sample, followed by post PCR clean-up and Nextera XT dual-indexed library preparation, according to the manufacturers’ instructions. Sequencing libraries were paired-end sequenced (2×150 bp) on an Illumina NextSeq 550 apparatus, as previously described[18]. Sequence read quality analysis and mapping was conducted using the bioinformatics pipeline implemented in INSaFLU (https://insaflu.insa.pt/; https://github.com/INSaFLU; https://insaflu.readthedocs.io/en/latest/; as of 10 March 2021), which is a web-based (and also locally installable) platform for amplicon-based next-generation sequencing data analysis[18]. We performed raw reads quality analysis using FastQC v0.11.9 (https://www.bioinformatics.babraham.ac.uk/projects/fastqc), followed by quality improvement using Trimmomatic v.0.27 (http://www.usadellab.org/cms/index.php?page=trimmomatic; HEADCROP:30 CROP:90 SLIDINGWINDOW:5:20 LEADING:3 TRAILING:3 MINLEN:35 TOPHRED33), with reads being conservatively cropped 30 bp at both ends for primer clipping. Reference-based mapping was performed against the Wuhan-Hu-1/2019 reference genome sequence (https://www.ncbi.nlm.nih.gov/nuccore/MN908947.3; NC_045512.2) using the Burrow-Wheeler Aligner (BWA_MEM) v.0.7.12 (r1039) (http://bio-bwa.sourceforge.net/)[19] integrated in multisoftware tool Snippy (https://github.com/tseemann/snippy) available in INSaFLU. The obtained median depth of coverage throughout the genome for CoV-2-D and CoV-2-G samples (except two samples excluded due to low coverage) was 4807 (IQR=3969-5242) and 5154 (IQR=4802-5439), respectively. Variant (SNP/indels) calling was performed over BAM files using LoFreq v.2.1.5 (*call* mode, including --*call-indels*)[20], with indel qualities being assessed using Dindel[21]. Mutation frequency analysis was dynamic and contingent on the depth of coverage of each processed site, e.g. minor mutations at “allele” frequencies of 10%, 2% and 1% (minimum cut-off used) were validated for sites with depth of coverage of at least 100-fold, 500-fold and 1000-fold, respectively. The median depth coverage per site for all validated mutations in CoV-2-D and CoV-2-G samples was 4219 (IQR=2508-6649) and 6424 (IQR=3076-10104), respectively. In order to assess if proximal SNPs and/or indels belong to the same mutational event (and thus, avoid overestimating the mutation rate), we identified all consecutive mutations separated by ≤12 bp. The mutations more likely to represent a single mutation event, i.e., those with similar frequencies (differing by ≤ 2.5%), were further visually inspected using IGV (http://software.broadinstitute.org/software/igv/) to confirm/exclude their co-localization in the same reads. In total, this curation led to the identification 37 SNPs/indels that were collapsed into 13 complex or multi-nucleotide polymorphisms (MNP). The effect of mutations on genes and predicted protein sequences was determined using Ensembl Variant Effect Predictor (VEP) version 103.1 (https://github.com/Ensembl/ensembl-vep; available as a self-contained Docker image)[22]. To obtain a refined annotation including all ORF1ab sub-peptides, the GFF3 genome annotation file (relative to the reference Wuhan-Hu-1/2019 genome of SARS-CoV-2, acc. no. NC_045512.2) available in the coronapp COVID-19 genome annotator (http://giorgilab.unibo.it/coronannotator/)[23]was adapted to generate an annotation GTF file for input for the --*gtf* parameter. The parameter --*distance* was set to 0. **Supplementary Table 1** summarizes all mutations detected in this study and their distribution across clinical, ancestral cultures and end-point cultured lines (15^th^ passage). SARS-CoV-2 consensus sequences obtained directly from clinical samples for CoV-2-D (Portugal/PT0054/2020) and CoV-2-G (Portugal/PT1136/2020) viruses are available in GISAID under the accession numbers EPI_ISL_421457 and EPI_ISL_511683, respectively. Reads generated at the end of the experimental evolution study were deposited in the European Nucleotide Archive (ENA) (https://www.ebi.ac.uk/ena/data/view/PRJEB43731).

### Simulations of the neutral mutation accumulation

To obtain a non-equilibrium neutral expectation of the site frequency spectrum of mutations, we performed forward-simulations to model mutation accumulation using the mutation rate inferred from the experiment. We model an organism with a bi-allelic genome of size L=30000 (~SARS-CoV-2). An initially isogenic population undergoes 15 cycles of growth, mutation and bottleneck, according to the following life cycle:

1. A clonal population starts with an inoculum size of 2000.
2. Each genome replicates X times. We assume the burst size X to be Poisson distributed with mean 1000.
3. For each of the replicating genomes we introduce a Poisson number of mutations with mean 0.1 (corresponding to a rate of 3.3×10^-6^ nt^-1^ cycle^-1^). We assume mutations to emerge with uniform probability in the parental genome and we allow for back-mutation.
4. After replication and mutation, we sample 1/1000 of the individual genomes.
5. Repeat steps 2-4, 15 times.

We validated the simulation code by confirming expected outcomes: mutations accumulate linearly over time and the posterior estimation of the mutation rate retrieves the original value (bottom plot in **Fig. S3b**).

After 15 cycles we collect the artificial genomes from 100 independent simulations, and compute their site frequency spectrum as in the experiment. In order to test whether cross-well contamination could justify the observed site frequency distribution, we modified the previous algorithm by introducing migration. At each cycle t, after each bottleneck event, a fraction of viral genomes (m=0.1) is replaced by migrants sampled from a pool of genomes that have undergone *t* cycles of growth. The algorithm was written in R (version 3.6.1) and the results analyzed in RStudio[24].

### Mutation accumulation rates in all, synonymous and non-synonymous sites

To quantify the rate at which mutations accumulate during the experiment, we compute 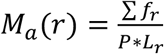, where *f_r_* is the frequency of all mutations observed in region *r*, P=15 the number of passages and *L_r_* the length of region *r*. For the genome-wise mutation accumulation *L_r_* = 29903, the entire genome of SARS-CoV-2. We also computed the mutation accumulation rates at synonymous and non-synonymous sites. In these cases, the synonymous rate is given by 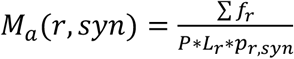, where *p_r,syn_* is the proportion of mutations in region *r*, leading to synonymous changes. Equivalently the non-synonymous rate is 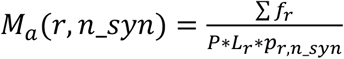. In practice, assume the region of interest has sequence *r*: ATGTTT. For each base we count the proportion of mutations that would change (or not) the corresponding amino acid. In the example *p_r,n_syn_* = 3/3 + 3/3 + 3/3 + 3/3 + 3/3 + 2/3 = 17/18, 17 mutations out of the possible 18 are non-synonymous and only one is synonymous (ATGTTc). Therefore, in this example, the total size is *L_r_* = 6, *P_r,n_syn_* = 17/18 and *p_r,syn_* = 1/18. Following this method, we calculated the genome-wise and gene-specific mutation accumulation rates in all, synonymous or non-synonymous sites (**Fig. 1-4** and **Fig. S4-5**). The genomic sequences of each region were retrieved from NCBI (entry: <u>NC 045512</u>).

**Fig. 1.**
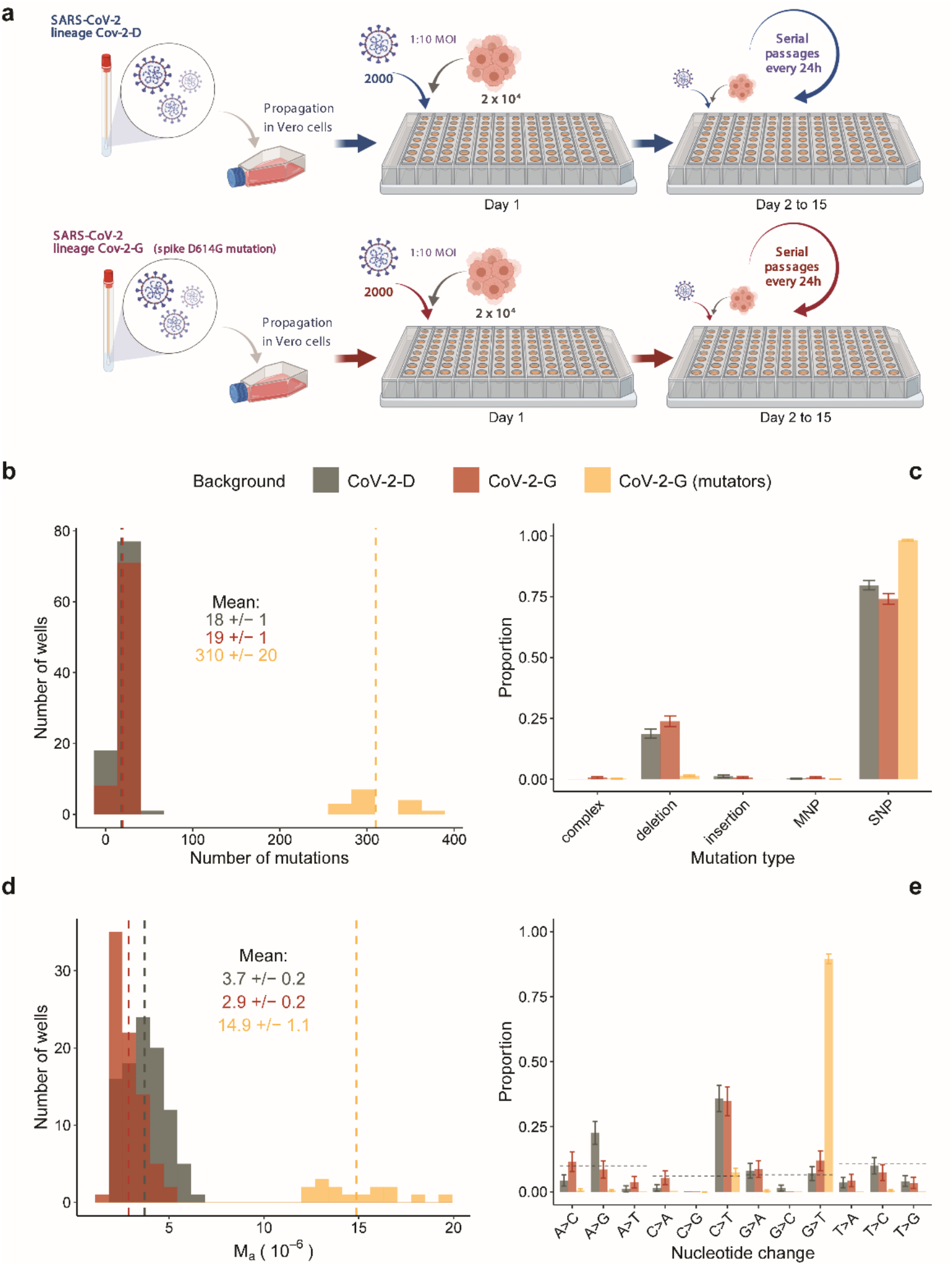
Experimental design and mutation accumulation after 15 passages of SARS-CoV-2 evolution. **a,** Schematic of the experimental design of the mutation accumulation experiments where two viral backgrounds were propagated in Vero cells (figure created with BioRender.com). **b,** Number of mutations observed in each well and group. 15 lines of the CoV-2-G background accumulated a larger number of mutations and thus were defined as mutators (gold). The means of each group are presented by vertical dashed lines and reported in the figure (+/− 2SEM). **c,** Proportion of mutation types in each group. Complex mutations and multi-nucleotide polymorphisms (MNP) are defined in the *Methodology*. **d,** Mutation accumulation per base per infection cycle (*M_a_*) was calculated by summing the observed mutation frequencies as: 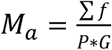, where *P* is the number of passages (*P*=15) and G is the SARS-CoV-2 genome length (G=29903). The means of each group are presented by vertical dashed lines and reported in the figure (+/− 2SEM). **e,** Proportion of observed nucleotide changes. Dashed lines indicate the expectation given the genome composition under equal mutation probability for each type of nucleotide change. Vertical bars in panels **c** and **e** represent the 95% confidence interval computed as 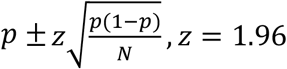,*z* = 1.96.

**Fig. 2.**
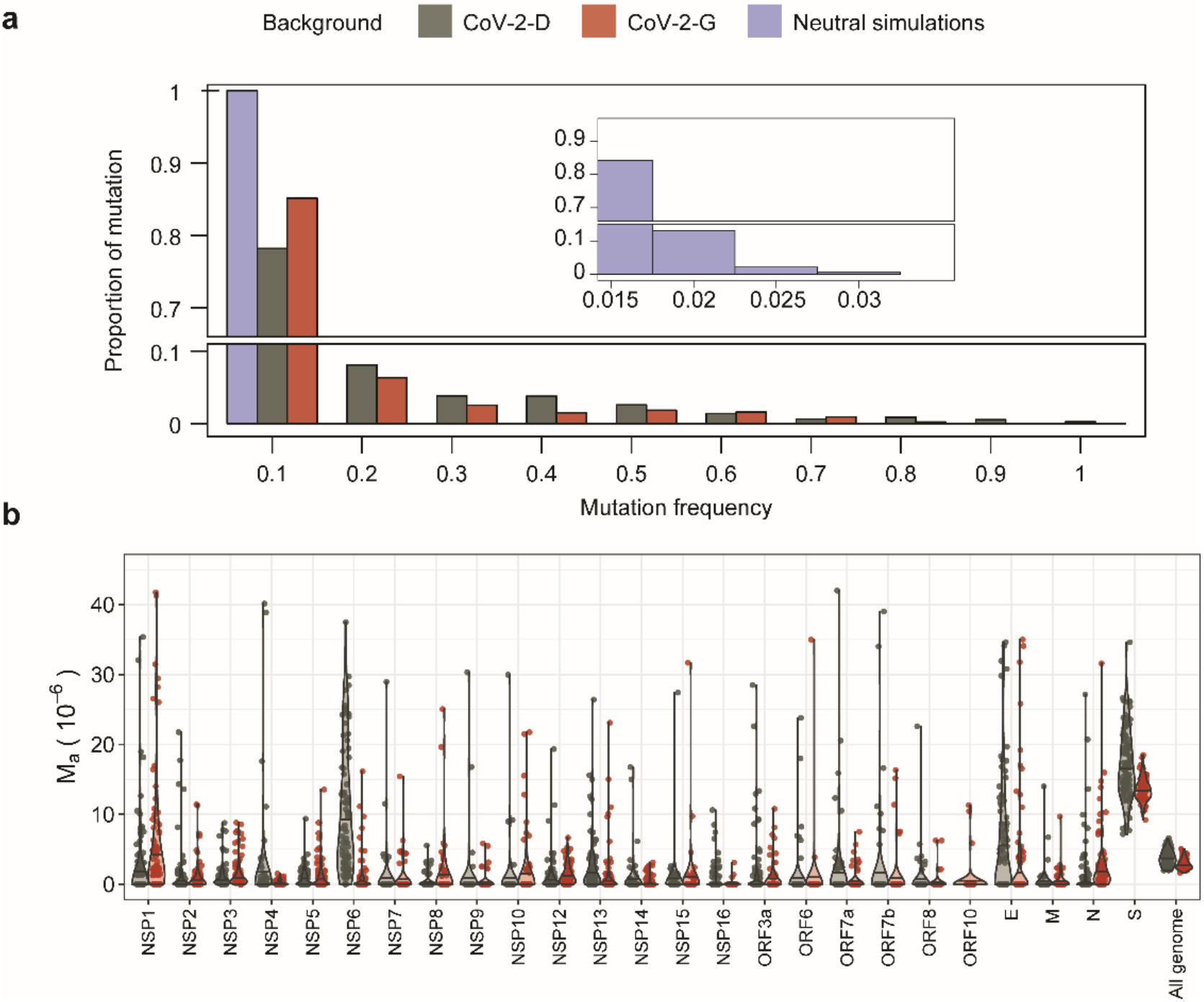
Site frequency spectrum and heterogeneity across genes. **a,** Proportion of mutations with a given frequency after 15 cycles of propagation in the CoV-2-D and CoV-2-G genetic background or under a simulated neutral model of mutation accumulation. The bump observed at high frequencies in the data is not compatible with the expectation of the neutral model. **b,** Per-base mutation accumulation (*M_a_*) computed for each gene and for the entire genome shows heterogeneity. The spike gene has the largest accumulation rate in both backgrounds (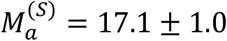, 13.5 ± 0.4 · 10^-6^, for the CoV-2-D and CoV-2-G respectively), which is more than 4 times their genomic average. For resolution purposes, few outliers with *M_a_* above 45 are not shown (see full set in **Fig. S4**).

**Fig. 3.**
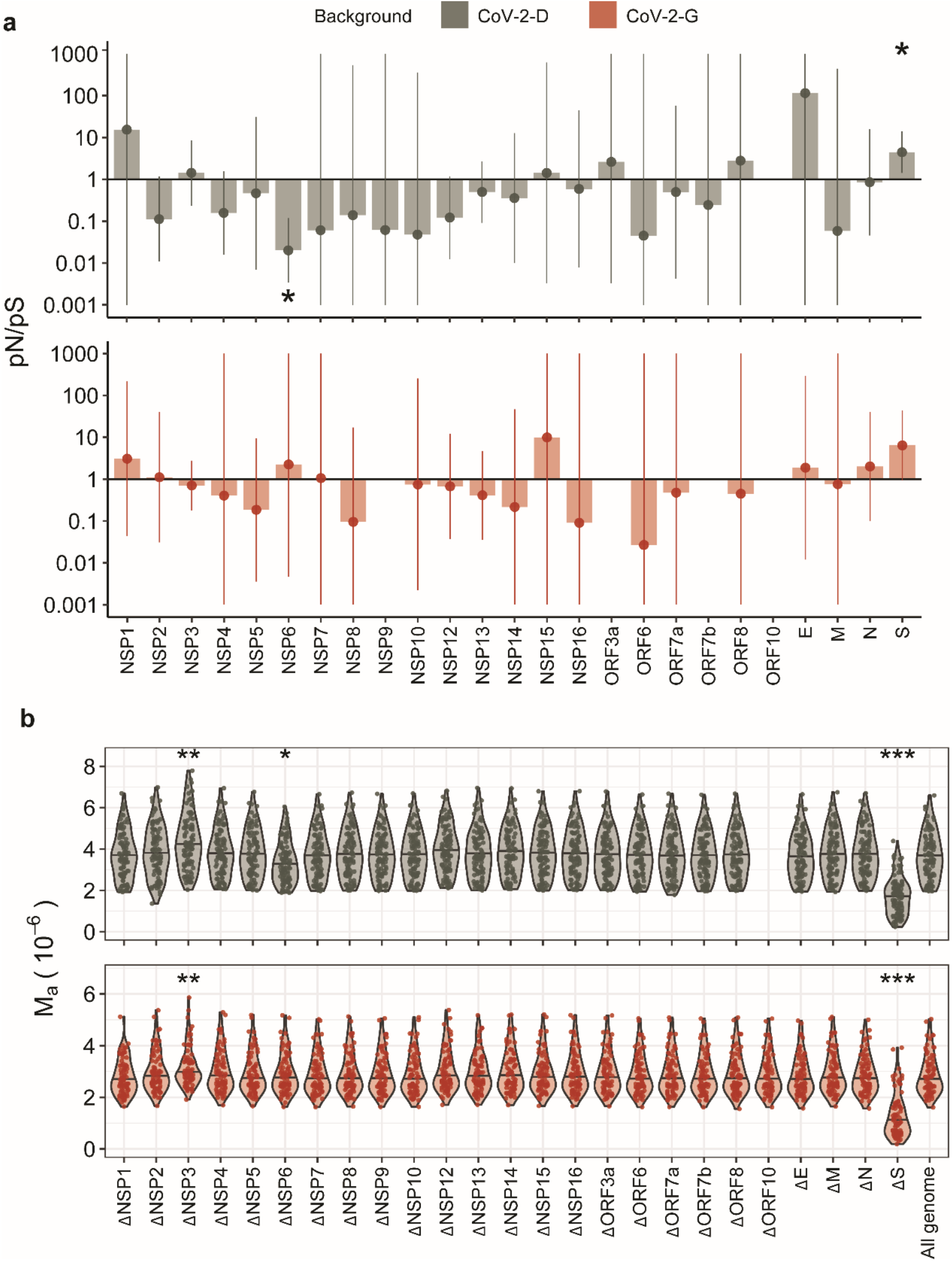
Gene-wise signs of selection. **a,** The relative proportion of non-synonymous to synonymous polymorphism, *pN/pS*, was computed for each gene and genetic background (see *Methodology*). The horizontal line indicates the expectation under neutrality (pN/pS=*1*), values above suggest positive selection while values below suggest purifying selection. Vertical bars show the 95% confidence intervals and the stars indicate the genes where such interval does not include 1. For the sake of resolution, we show the confidence intervals within the [10^−3^,10^3^] range. **b,** Identifying the genes that affect the estimation of mutation rate. Per-base mutation accumulation (*M_a_*) was computed for the entire genome or by excluding each gene on at the time (e.g. ΔS). The stars indicate the cases where removing the gene leads to an estimation of *M_a_* significantly different from the all genome (non-parametric Wilcox test, p-value < 0.05 (*), 0.01 (**) or 0.001 (***)).

**Fig. 4.**
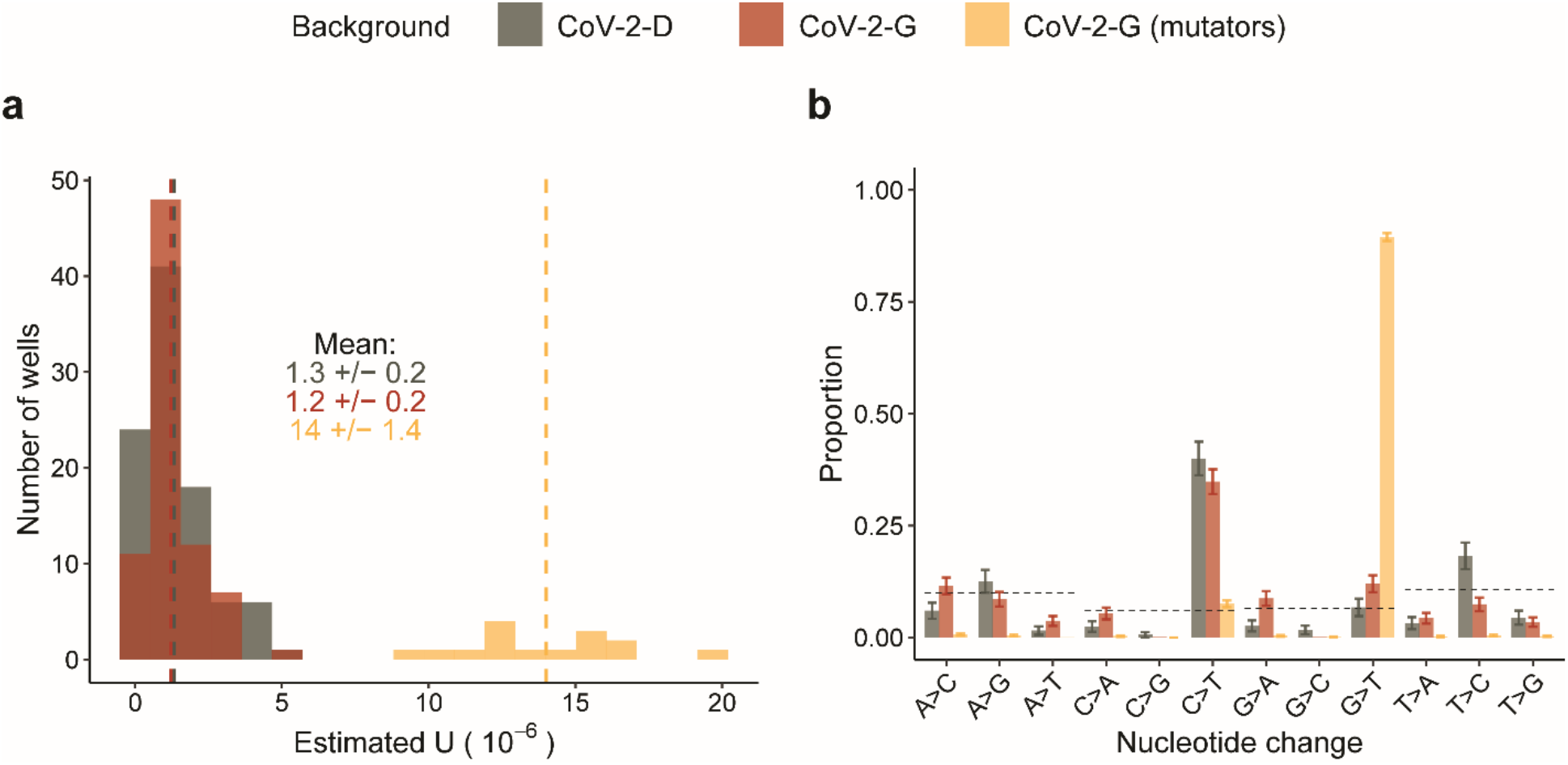
Estimation of mutation rates and bias excluding outlier genes. **a,** The per-base per-infection cycle mutation rate was calculated by summing the observed mutation frequencies as: 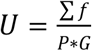, where *P* is the number of passages (*P*=15) and G is the length of SARS-CoV-2 genome excluding the Nsp3, Nsp6 and Spike genes (29903-5835-870-3822=19376). The means of each group are presented as vertical dashed lines and reported in the figure (+/− 2SEM). **b,** Proportion of nucleotide changes observed excluding the Nsp3, Nsp6 and Spike genes. Dotted lines indicate the expectation given the genome composition under equal mutation probability for each type of nucleotide change. Vertical bars represent the 95% confidence interval computed as 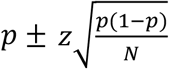, *z* = 1.96.

### pN/pS calculation and confidence interval

Within a given region *r*, we computed *pN(r)* as the summed frequencies of all the observed non-synonymous mutations over the number of all possible non-synonymous changes in that region: 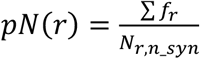, where *N_r,syn_ = 3L_r_p_r,n_syn__*. Equivalently, we computed the synonymous counterpart: 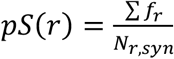. In the previous example, within the region *r*: ATGTTT, *N_r,n_syn_* = 17 while *N_r,syn_* = 1. Finally, the *pN/pS* statistics is the ratio of *pN* and *pS* and its expected value is 1 under neutrality. To test the deviation from 1, we report the *pN/pS* together with its confidence interval. Being the *pN/pS* a ratio of proportions, we computed the 95% confidence intervals of risk ratios, specifically as 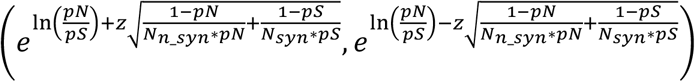 with critical *z* = 1.96.

## Results

### Experimental evolution design and ancestor backgrounds

Two SARS-CoV-2 viral strains were isolated from two non-related patients for continuous propagation in Vero cells (see Methodology, **Fig. 1a**). These were chosen according to their polymorphism at amino acid position 614 of the spike protein: CoV-2-D carries a D and CoV-2-G carries a derived mutation which changes the D into a G. This D614G mutation in spike emerged early in the pandemic, increased the infectivity of the virus and became prevalent worldwide[25]. Here, we want to test for differences in their mutation rates, spectrum and/or in the selective forces as the strains are propagated in cells.

In order to discriminate *de novo* mutations from standing genetic variation, we identified the mutations (relative to the Wuhan-Hu-1/2019 reference genome sequence, Wu et al., 2020) that were already present at the start of our evolution experiment (see the list and their frequencies in **Fig. S1**).

If the mutation rate is similar to that of the mouse hepatitis virus (MHV) or that of the SARS-CoV (about 3.5×10^-6^ and 2.5×10^-6^ nt^-1^ cycle^-1^, respectively)[5,7] hundreds of mutations should accumulate, many of which are expected to be neutral but some could reflect adaptation to the experimental conditions.

### Mutation accumulation and spectrum after 15 passages of SARS-CoV-2 evolution

We considered *de novo* mutations those that reached a frequency of at least 1%, supported by a minimum of 10 reads and that were not detected in either the ancestor or the original clinical isolate from which the ancestor was derived (full list in **Supplementary Table 1**). Propagation of the 96 CoV-2-D derived lines resulted in 1753 *de novo* mutations, while the 96 lines derived from CoV-2-G resulted in 6181 *de novo* mutations (n=94 as in two lines the sequencing had poor coverage) (**Fig. 1b**). The much higher number of mutations in the CoV-2-G background, compared to CoV-2-D, is explained by 15 of these lines where many more mutations were observed (**Fig. 1b**). These lines, hereafter referred to as mutators, are characterized by a larger proportion of SNPs compared to the non-mutator lines where, instead, deletions account for more than 20% of all *de novo* mutations (**Fig. 1c**).

The frequency of mutator clones was estimated to be between 1 and 2% after 15 infection cycles, since these were the frequencies measured for the vast majority of mutations observed in the mutator lines. The genetic cause of the mutator phenotype is difficult to determine but it could likely be hidden within the mutations that occurred in the RNA-dependent RNA polymerase (Nsp12) and/or in the error-correcting exonuclease protein (Nsp14)[8]. Indeed, looking at the mutations that are specific to the lineages with mutators, we found 8 non-synonymous mutations in Nsp12 (one leading to a stop at amino-acid 670) and 9 non-synonymous mutations in Nsp14 (one leading to a stop at amino acid 78) (**Supplementary Tables 2-3**). Any of these mutations could potentially lead to the observed change in mutation rate, but none of these have been associated with an increased mutational load of the circulating viruses[27].

Next, we obtained the per-base per-passage rate at which mutations accumulated (Ma), from the frequencies of the observed mutations. As a 24 h passage in our experiment corresponds to ~1 cell replication cycle (see *Methodology*), we hereafter report such rate of mutation accumulation per unit of replication cycle (nt^-1^ cycle^-1^). Interestingly, the non-mutator lines of CoV-2-G show a significantly lower accumulation rate compared to the CoV-2-D lines (P<10^-6^, Two-sample Kolmogorov-Smirnov test) (**Fig. 1d**). However, this difference between the two backgrounds is more likely due to differences in selection rather than differences in mutation rates, as we will explain later on.

The SNPs accumulated over 15 passages show that both genomic backgrounds have a strong propensity to accumulate C->T mutations (**Fig. 1e**), a well-known bias of SARS-CoV-2[28]. In the mutator lines, the main mutation bias changed from C->T to G->T (**Fig. 1e**), also observed in SARS-CoV-2 samples collected during the recent COVID-19 pandemic[29,30].

It is important to notice that, both the accumulation of mutations and the biases we observe in the data (**Fig. 1b-e**) might have been shaped by selection and deviate from a neutral rate and spectrum of mutations. In fact, on one hand positive selection can increase the frequencies of beneficial mutations and on the other hand purifying selection can purge the deleterious alleles. Therefore, we next looked for evidences of selection in the mutation accumulation data.

### Signs of selection: Site frequency spectrum and heterogeneity across genes

In serial propagation experiments with SARS-CoV-2, it is extremely difficult to pick a single virus[31]. In our experiment the effective population size is considerably large (see *Methodology*), and thus could be insufficient to remove the effects of either positive or negative selection[10]. Indeed, several patterns in the data indicate that selection played a significant role in the experimentally evolved SARS-CoV-2 lines.

The distribution of allele frequencies in a sample, *i.e*. the site frequency spectrum, has a well-known theoretical expectation under a simple equilibrium neutral model of molecular evolution (Chap. 5 pg. 233 of B. Charlesworth & D. Charlesworth., 2010). But, this distribution is sensitive to the action of selection and also to complex demographic events, such as population bottlenecks. Given the bottlenecks occurring in our experiments and the slow evolutionary time elapsed during the 15 infection cycles, the neutral theoretical expectation at equilibrium may not apply. To obtain a non-equilibrium expectation of the site frequency spectrum, we performed forward-simulations (see *Methodology*). We assumed that neutral mutations occur at a rate of 3.3×10^-6^ nt^-1^ cycle^-1^, similar to that estimated from the data, and simulated populations evolving under neutrality. The site frequency spectrum of the mutations accumulated in both CoV-2-D or CoV-2-G lines deviates significantly from the neutral expectation predicted by the simulations (**Fig. 2a**). High frequency mutations are not expected under neutrality (mutations with frequencies above 30% are reported in **Fig. S2**). To test whether possible contamination among wells could explain the observed site frequency spectrum, we performed additional simulations with migration (see *Methodology*). Even when considering a migration rate of 10%, the neutral site frequency spectrum is still incompatible with the experimental data (**Fig. S3a**). Furthermore, the 10% migration between wells should not significantly change the estimation of mutation rate (**Fig. S3b**). Thus, the data strongly suggest that positive selection has increased the frequency of beneficial mutations and skewed the spectrum of the mutations (**Fig. 2a**).

A second evidence of selection comes from the considerable variation in the rate of mutation accumulation observed across the SARS-CoV-2 genome (**Fig. 2b, Fig. S4**). When excluding the mutator lines, the S gene, which codes for the spike protein, has the highest rate of mutation accumulation among the different genes (**Fig. 2b**). Remarkably, the spike accumulated 13.5±0.4×10^-6^ nt^-1^/cycle^-1^ mutations in the CoV-2-G genotype (excluding mutators), and 17.1±1.0×10^-6^ in the CoV-2-D genotype, about five-fold the corresponding genomic averages, suggesting the strong action of positive selection. In the mutator lines, the spike gene evolved ~2 times faster than the non-mutators (**Fig. S4**). This observation is in contrast with the expectation of a constant increase in mutation rate across the genome and suggests that more complex selective forces might be acting on the mutator phenotype (see the heterogeneity of the mutation rate across the CoV-2-G mutator genome in **Fig. S4**). Overall, the data confirmed that selection has shaped the way mutations accumulated. Therefore, in order to obtain a more accurate quantification of the spontaneous rate of mutation, we performed a more systematic analysis of the sites under selection.

### Identifying regions under selection

From the frequencies of all mutations observed in the CoV-2-D and CoV-2-G non-mutator lines, we computed an accumulation rate of 3.7×10^-6^ and 2.9×10^-6^ nt^-1^ cycle^-1^, respectively (**Fig. 1d**). Given that during our experiment, selection affected the allele frequencies (**Fig. 2**), such rates may deviate from the spontaneous mutation rates of the virus. In order to attempt to estimate the spontaneous mutation rate we first focused on synonymous mutations, which, if neutral, should accumulate at the rate at which they occur[33]. Focusing on the synonymous changes, we estimated a basic mutation rate of 3.8×10^-6^ nt^-1^ cycle^-1^ for the CoV-2-D background and 1.2×10^-6^ nt^-1^ cycle^-1^ for the CoV-2-G (**Fig. S5a**). However, the rate of non-synonymous mutation in CoV-2-D is lower than the synonymous one (**Fig. S5a-b**), suggesting the action of purifying selection on non-synonymous sites or positive selection on the synonymous sites, leading to their increase in frequency[34,35]. To distinguish between the two cases, we compared the accumulation rate of synonymous mutations in the entire genome 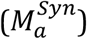, with the accumulation rate of synonymous mutations excluding one gene at the time 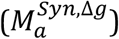. This approach revealed that a remarkable accumulation of synonymous mutations in the Nsp6 gene led to the overestimation of the mutation rate in the CoV-2-D background (**Fig. S5c**). In contrast, for the CoV-2-G background this approach indicates that the estimation of 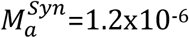 nt^-1^ cycle^-1^ is homogeneous across the genome and can provide a first estimation of its mutation rate (**Fig. S5d**).

Since the synonymous mutations alone could not provide a correct estimation of mutation rate, we followed a different approach: identify the regions under selection in either the CoV-2-D or CoV-2-G lines and exclude them from the estimation of the spontaneous mutation rate. First, we compared the relative accumulation of non-synonymous and synonymous mutations, via the *pN/pS* statistics (equivalent of *dN/dS* for polymorphic samples, see *Methodology*). In the CoV-2-D background, the *pN/pS* of the S and the Nsp6 genes significantly differ from 1 (**Fig. 3a**). The spike accumulated more non-synonymous mutations consistent with the action of positive selection (*pN/pS*=4.4, 95% confidence interval: [1.4,13.9]), while the Nsp6 accumulated more synonymous mutations, consistent with our previous findings (*pN/pS=0.02, 95%* confidence interval: [0.00,0.12], **Fig. S5c**). In particular, the synonymous change A11041G was found in 88 evolved populations (out of 96), but also at frequency below our 1% threshold in the ancestral population, suggesting that such mutation was incorrectly considered as *de novo* and that the estimated mutation accumulation in Nsp6 was the resulting artifact.

Due to the limited number of mutations within each gene and the fact that we are comparing evolving populations (rather than divergent species), the *pN/pS* may lack the power to identify additional regions under selection[36]. To overcome this issue and to identify additional genes affecting the estimation of the mutation rate, we computed the rate of mutation accumulation excluding one gene at the time and compared this with the entire genome (see *Methodology* and **Fig. 3c-d**). With this outlier-detecting method we confirmed that the S and the Nsp6 genes affected the estimation of mutation rate in the CoV-2-D strain, we could observe that the S gene is likely under selection also in the CoV-2-G strain, and we identified Nsp3 as an additional region with a different rate of mutation accumulation (**Fig. 3c-d**). In particular, Nsp3 accumulated fewer mutations than the genomic average in both CoV-2-D and CoV-2-G strains, suggesting the action of purifying selection (**Fig. 3c-d**).

Overall, we conclude that during our experiment, the spike protein was under strong selection in both backgrounds, but also other genes biased the estimation of mutation rate.

### Estimation of mutation rates and bias excluding genes with signs of selection

Non-neutral processes have shaped the allele dynamics in our experiment. To get a more realistic estimate of the mutation rate prior to selection, we excluded from the analysis the Nsp3, Nsp6 and S genes, which have shown signs of selection in at least one of the two backgrounds (**Fig. 3**). By doing this, we estimate a spontaneous mutation rate of 1.3±0.2×10^-6^ nt^-1^ cycle^-1^ for the CoV-2-D background and 1.2±0.2×10^-6^ nt^-1^ cycle^-1^ for the CoV-2-G (excluding mutators) (**Fig. 4a)**. The estimated mutation rate is similar across backgrounds, suggesting that the previously observed differences were due to selection (**Fig. 1d** and **S5**). Importantly, the estimated mutation rate of CoV-2-G is consistent with that obtained using the synonymous mutations only (see **Fig. S5a**) and the estimated mutation rate of CoV-2-D is consistent with that obtained using the synonymous mutations only and excluding the Nsp6 (see **Fig. S5c**). We quantified again the relative proportion of single nucleotide changes and confirmed that both backgrounds have a spontaneous bias towards C>T mutations and the mutator changes this bias towards G>T mutations (**Fig. 4b**).

Excluding the genes with signs of selection was our best attempt at quantifying the spontaneous mutation rate of SARS-CoV-2. However, it is important to note that this may still underestimate the real one due to the fact that we ignored mutations with a frequency below the 1% threshold.

### Convergent targets of selection on Spike

The spike protein showed clear signs of adaptation during our evolution experiment, so we next focused on the specific sites under selection and compared them with the new spike variants that spread in the human population. We first quantified the level of convergence at the nucleotide and amino levels between CoV-2-D and CoV-2-G. We note that convergence between the two backgrounds reflects true independent origin of the mutations, as they were propagated and processed for sequencing independently. In contrast, convergence within replicates of the same background could also result from some possible cross-contamination or from undetected standing variation. At the amino level, 20 specific sites and 3 regions were hit independently in both backgrounds (**Fig. 5a, Supplementary Table 4).** We find high evolutionary convergence at the S1/S2 cleavage site: three distinct deletions (675-QTQTN-679 del; 679-NSPRRAR-685 del and 679-NSPRRARSVA-688) emerge multiple times in both backgrounds. Such changes have been previously shown to emerge rapidly in Vero cells and to be important for the virus cell tropism[37]. Apart from these deletions, mutations of the Arginine 682 were also highly convergent, most likely because they trigger a similar functional effect, i.e., knock out of the furin cleavage site[38]. Notably, another deletion in this region (678-TNSPRRARS-686 del) was frequently observed, still it was exclusive of CoV-2-D lines (*n*= 58), suggesting that the conformation changes mediated by D614G may influence the directionality of the evolution towards the knock out of the furin-cleavage site[39].

**Fig. 5.**
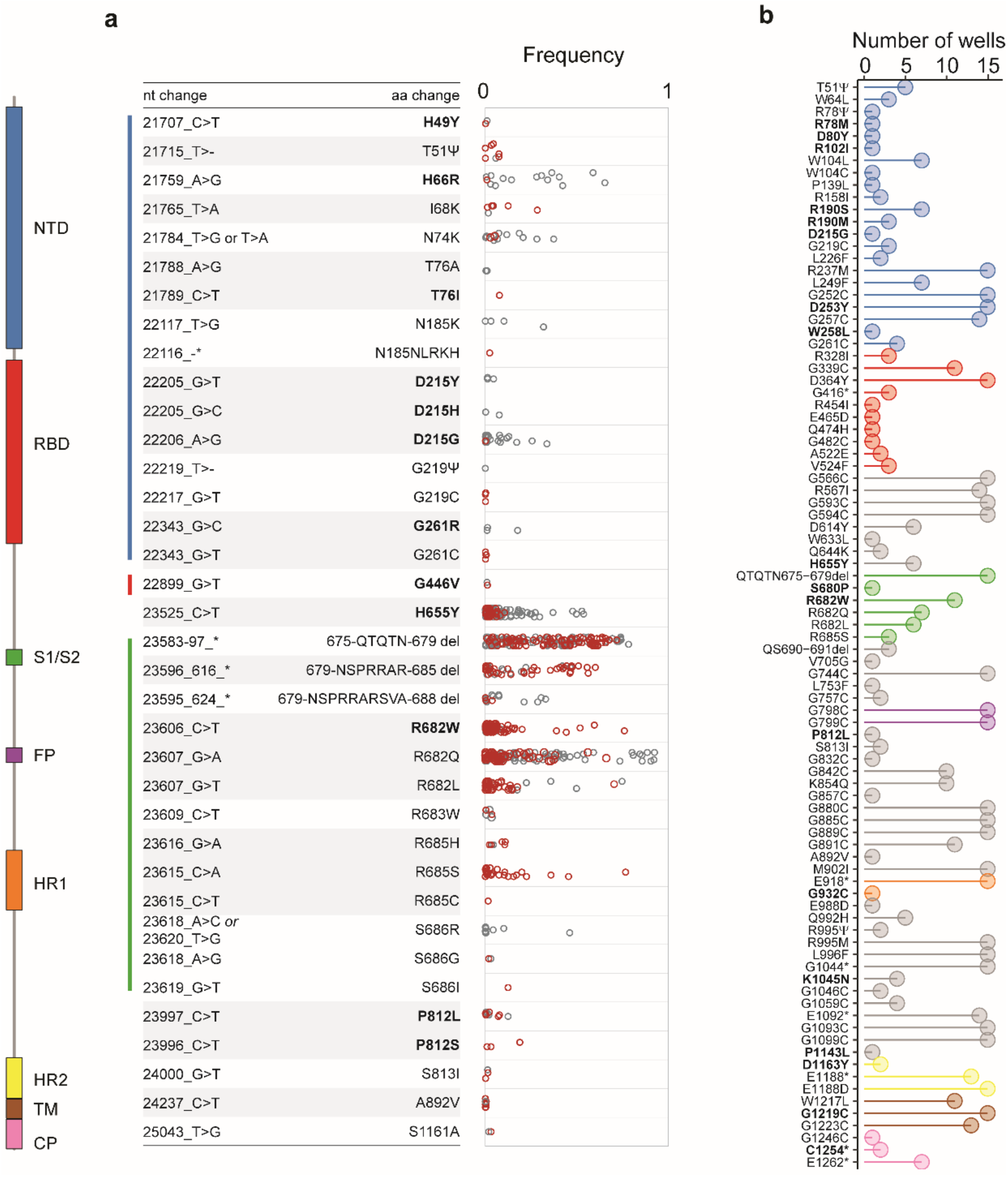
Convergent evolution in the Spike gene. **a,** Mutations on S observed in both CoV-2-D and CoV-2-G backgrounds and their frequencies in each well (open circles). **b,** Non-synonymous mutations on the spike detected in the populations where the mutators were observed (number of wells on the X-axis). The color annotation represents the N-terminal domain (NTD, 14–305), the receptor-binding domain (RBD, 319–541), the cleavage site (S1/S2, 669-688), the fusion peptide (FP, 788–806), the heptapeptide repeat sequences (HR1, 912–984 and HR2, 1163–1213), the TM domain (1213–1237), and cytoplasm domain (CP, 1237–1273). Amino acids changes in bold were also observed in the human population (as of 24 October 2021; https://nextstrain.org/ncov/gisaid/global) (full list in **Supplementary Table 6**).

Some level of evolutionary convergence could also be found for the structural genes N, E and M suggesting that adaptation could also have occurred in these genes (**Supplementary Table 5**).

The inferred mutators in the CoV-2-G background also carry many mutations in the spike protein including in the receptor binding domain -RBD- (amino acid changes at positions 328, 339, 364, 416, 454, 465, 474, 479, 482, 522 and 524) and multi cleavage site regions (positions 798 and 799) (**Fig. 5b**).

When scrutinizing the list of non-synonymous mutations in the spike that emerged during our experiment in both backgrounds or in the mutator lines, we found 24 amino-acid changes that were also observed in the natural population of SARS-CoV-2 (until the 24th of October 2021; https://nextstrain.org/ncov/gisaid/global) (full list reported in **Supplementary Table 6** and highlighted in bold in **Fig. 5**). Among these, we observed the mutations H655Y (present in the variant of concern Gamma, lineage P.1, originated in Brazil), D215G (present in the variant of concern Beta, lineage B.1.351, firstly identified in South Africa) and D253G (found in lineage B.1.426, mostly detected in the US) (**Fig. 5b**)[40].

## Conclusions and implications

The SARS-CoV-2 beta-coronavirus, first observed in the Wuhan province of China[6], has infected at least 246 million people causing more than a 5 million toll of deaths in the human population (as of 2 November 2021; https://covid19.who.int/). Since it was first sequenced[26] the virus has been accumulating 0.44 substitutions per week at close to linear rate. Here we estimate its rate of spontaneous mutation to be of the order of 10^-6^ per base per cell infection, consistent with previous estimations in other coronaviruses[9]. New beneficial mutations did spread to high frequencies and considerable convergent evolution was detected across different genomic backgrounds. We also observe viral populations with an increased mutation rate emerging just within 15 days of propagation in cells. This suggests that the mutation rate of SARS-CoV-2 can increase without significant loss of viability (at least in the short run) and that strategies to reduce viral fitness using mutagens should be tested with precaution[41,42].

Overall the results show the remarkable ability of SARS-CoV-2 to adapt to new environments, in particular via convergent evolution of its spike protein in cells, and is fully consistent to its rapid adaptation to different hosts[43,44].

## Data and materials availability

SARS-CoV-2 consensus genome sequences obtained directly from clinical samples for CoV-2-D (Portugal/PT0054/2020) and CoV-2-G (Portugal/PT1136/2020) viruses are available in GISAID under the accession numbers EPI_ISL_421457 and EPI_ISL_511683, respectively. Reads generated throughout the experimental evolution in this study were deposited in the European Nucleotide Archive (ENA) (https://www.ebi.ac.uk/ena/data/view/PRJEB43731). The code for the neutral model was deposited on https://github.com/AmiconeM/neutralviralpassage.

## Acknowledgments

We would like to thank the personnel of the IGC Genomic Facility for their assistance.

## Author contributions

IG, MJA and JPG designed the project. MJA and LZZ performed the culture and RNA extraction experiments. VB and JI performed the pre-sequencing wet-lab procedures, bioinformatic analysis and data analysis. SD and LV performed and supervised the wet-lab sequencing procedures. IG, MA performed the data analysis and the simulations. MJA, LV and JPG provided materials and reagents. IG wrote the initial draft of manuscript. VB, MJA, MA and JI contributed equally to this work. All authors contributed in the final writing of the manuscript and gave final approval for publication.

## Ethical statement

The Portuguese NIH is authorized by the Portuguese Authorities’ (General-Directorate of Health and the Authority for Working Conditions) to handle and propagate Risk group 2 and 3 microorganisms. All culture procedures were performed inside a class III microbiological safety cabinet in a containment level 3 facility. This study is covered by the ethical approval issued by the Ethical Committee (“Comissão de Ética para a Saúde”) of the Portuguese National Institute of Health.

## Additional information

Supplementary information is available for this paper. Correspondence and requests for materials should be addressed to.

