## Supplementary for "Mutation rate of SARS-CoV-2 and emergence of mutators during experimental evolution"

### Supplementary Figures

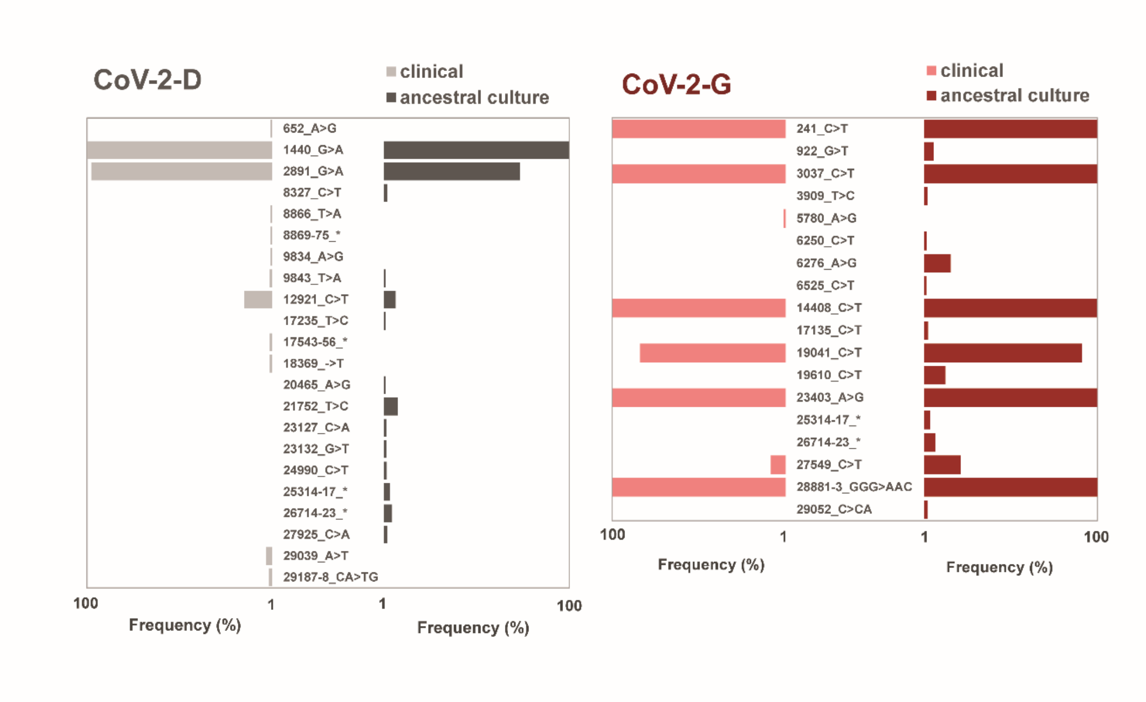


**Supplementary Fig. S1|** Mutations and their frequencies in the clinical isolates and their corresponding derived ancestors for the propagations. For the sake of figure simplicity, the complex mutations indicated in the figure with an asterisk correspond to: 8869-75_TTTGCCT>CAAACCA; 17543-56_TGTTCCTCGGAACT>AGTTCCGAGGAACA; 25314-17_GATC>TATG; 26714-23_TTTTGTGCTT>-GTTGTAC--.

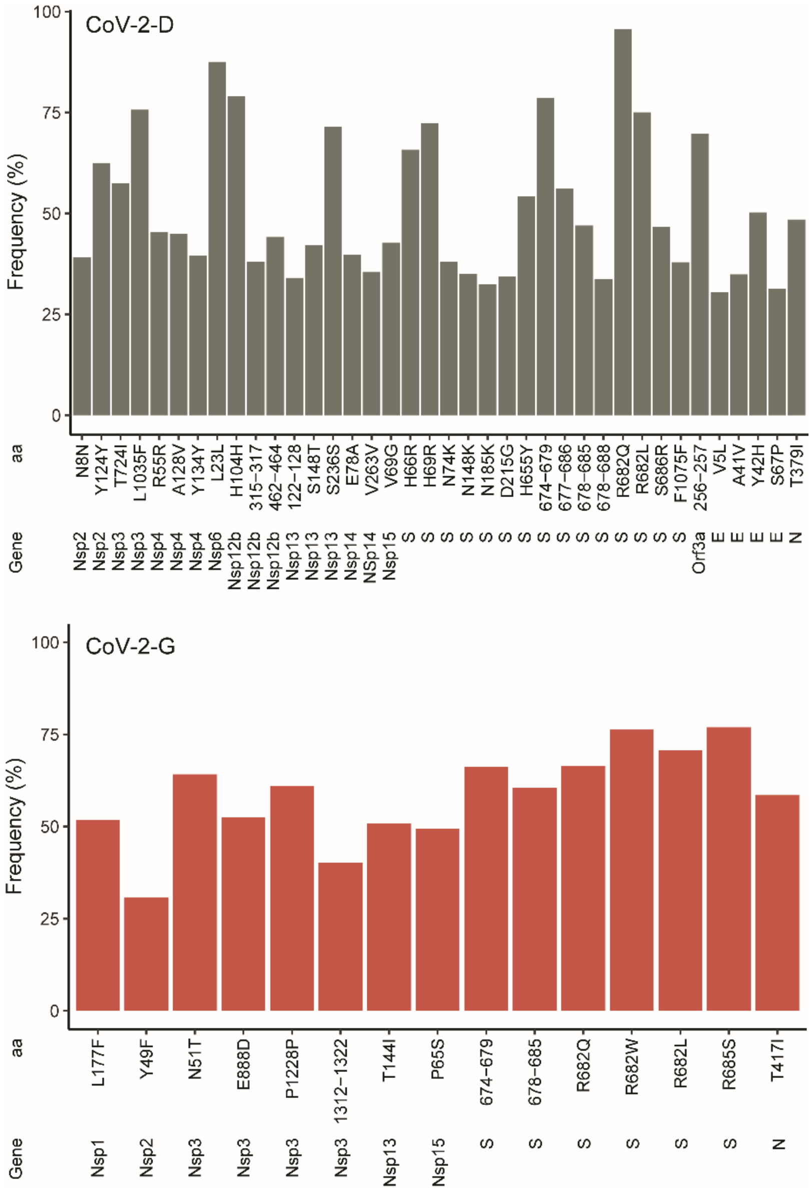


**Supplementary Fig. S2|** Mutations that reached frequency above 30% in each genomic background. The changes where deletions occur are indicated without letters (e.g. 674-679 corresponds to a deletion of YQTQTN to Y, see **Supplementary Table 1** for detailed annotation of all mutations).

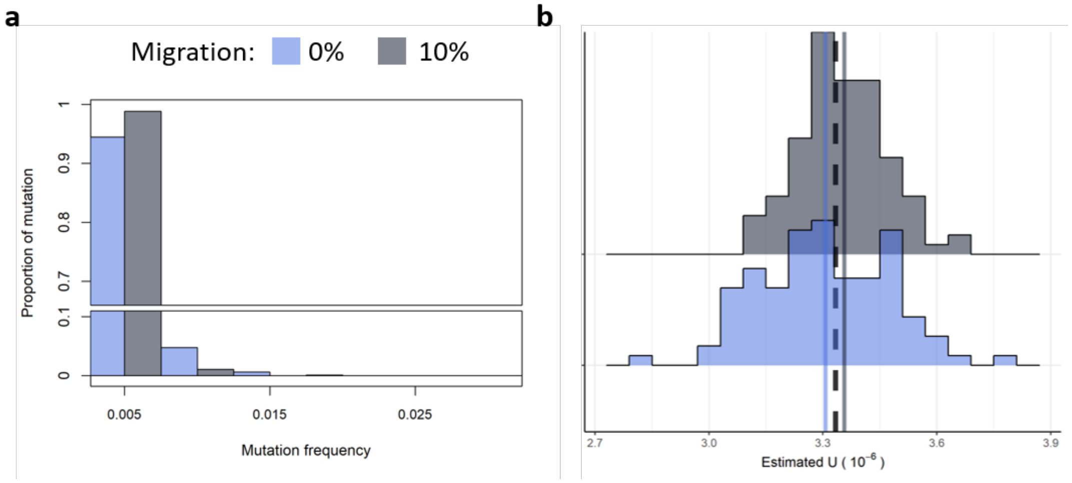


**Supplementary Fig. S3|** 10% cross contamination between wells cannot explain the frequency spectrum of the observed mutations nor can it affect the estimation of mutation rate. **a,** the neutral site frequency spectrum expected with or without 10% migration (or cross-contamination) between wells at each infection cycle. **b,** No effect of 10% migration on the estimation of mutation rate. The dashed line represents the simulated *U* while the continuous lines represent the average of the estimated *U* with or without cross.

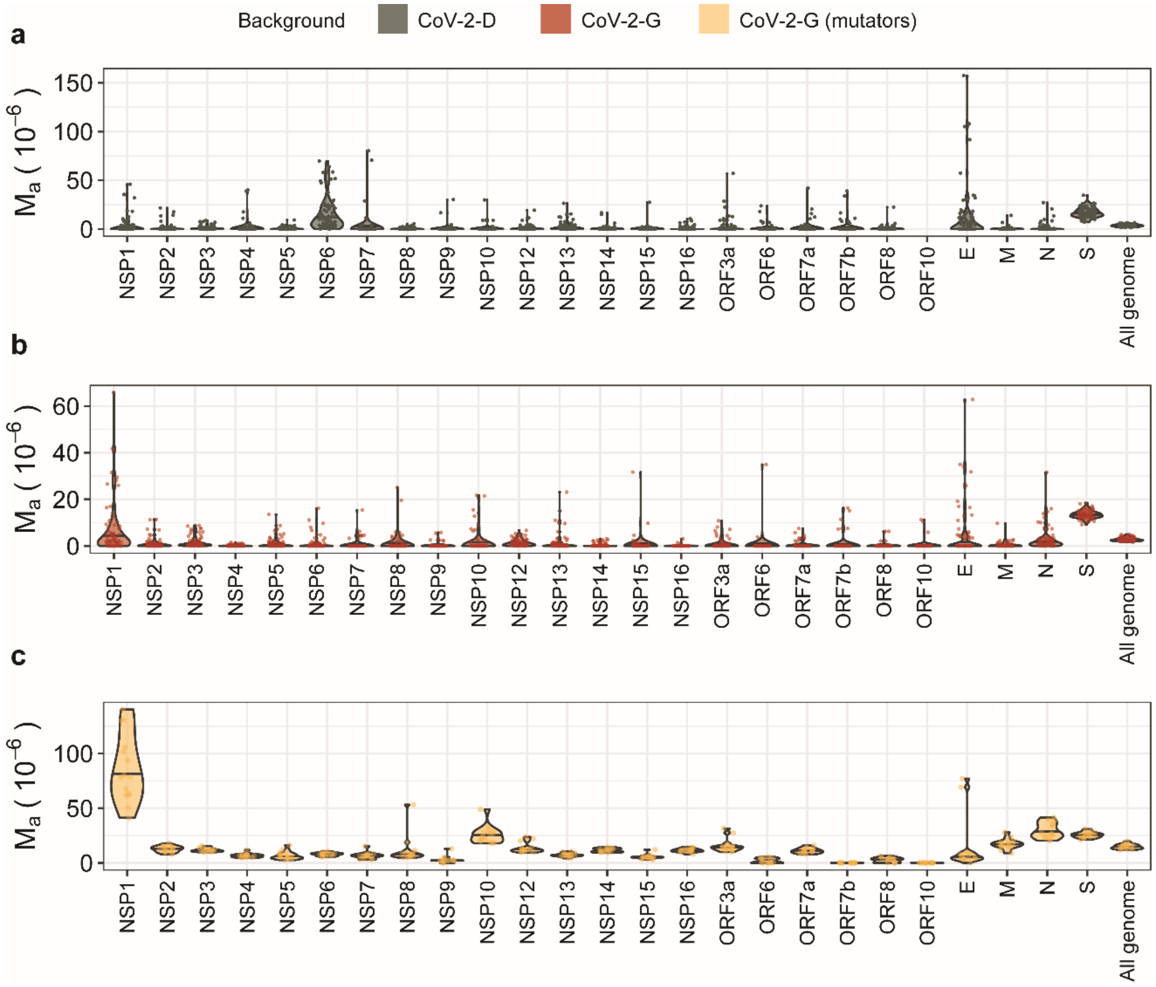


**Supplementary Fig. S4|** Heterogeneity of mutation accumulation across the genome of SARS-CoV-2. **a-c,** Per-base mutation accumulation ($M_a$) computed for each gene and for the entire genome in the CoV-2-D (n=96), CoV-2-G (non-mutators, n=79) and CoV-2-G (mutators, n=15) backgrounds, respectively.

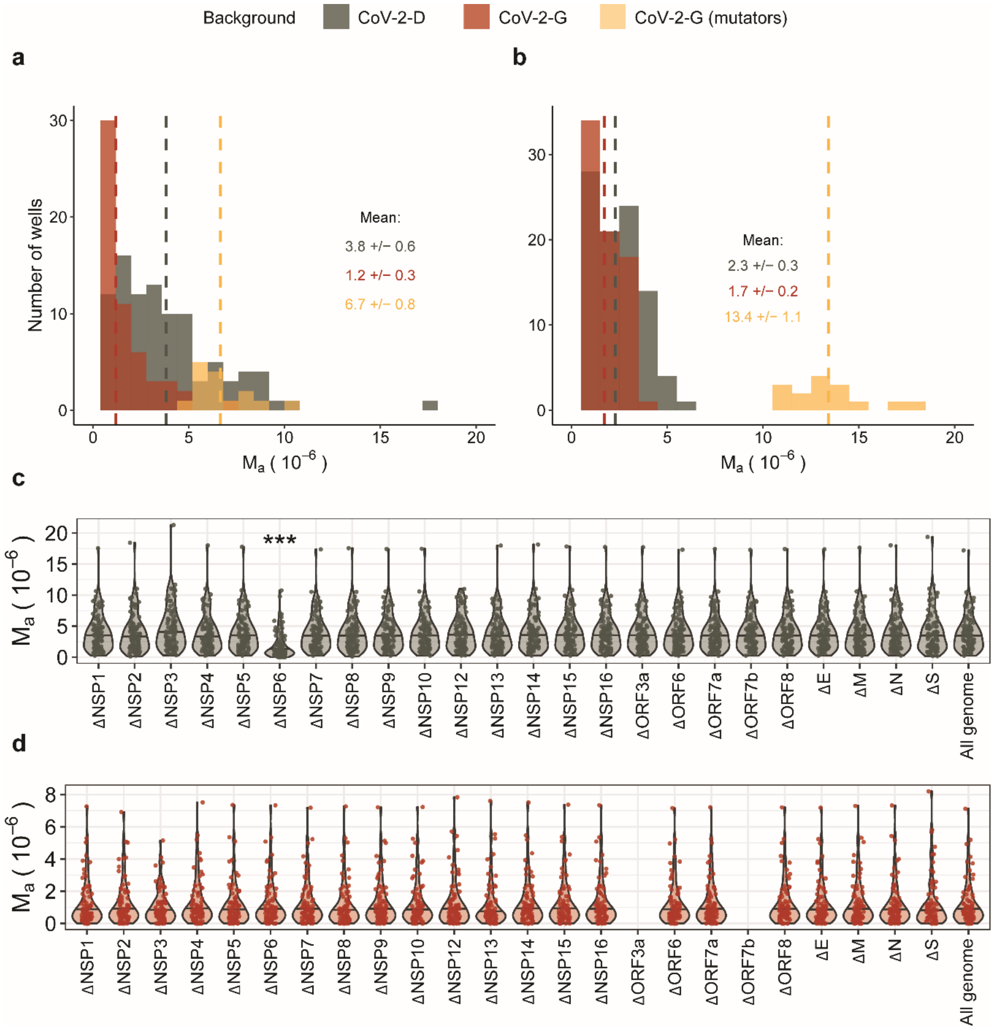


**Supplementary Fig. S5| a-b,** Per-base per-cycle mutation accumulation ($M_a$) in synonymous and non-synonymous sites. $M_a$ was calculated by summing the observed mutation frequencies as: $M_a = \frac{\sum f}{P*G}$, where $P$ is the number of passages ($P$=15) and G is the number of synonymous or non-synonymous sites in the genome of SARS-CoV-2 (G=6418 and 22846, respectively). The means of each group are presented by vertical dashed lines and reported in the figure (+/- 2SEM). **c-d,** The Nsp6 gene in CoV-2-D background is responsible for the overestimation of $M_a$ (panel **a**) and should be removed. Per-base mutation accumulation ($M_a$) was computed from the synonymous mutations throughout the entire genome or by excluding each gene on at the time (e.g. ΔS). The stars indicate the cases where removing the gene leads to an estimation of $M_a$ significantly different from the all genome (non-parametric Wilcox test, p-value < 0.05 (*), 0.01 (**) or 0.001 (***), after Benjamin-Hochberg correction).

**Supplementary Tables**

Supplementary Tables 1 to 6 are uploaded as Excel files.

**Supplementary Table 1| Mutation summary and accession numbers.** List of all detected mutations and their distribution across clinical, ancestral cultures and end-point cultured lines (15th passage). MNP = multi-nucleotide polymorphism; complex = mutation event comprising SNP and indels; * = stop codon; fs = frameshift. Sheet2 contains the European Nucleotide Archive (ENA) accession numbers for the read sequencing data.

**Supplementary Table 2| Mutator-specific mutations on the Nsp12 gene.** List of all mutations in the Nsp12 gene, unique to the mutator lines that emerged in the CoV-2-G background.

**Supplementary Table 3| Mutator-specific mutations on the Nsp14 gene.** List of all mutations in the Nsp14 gene, unique to the mutator lines that emerged in the CoV-2-G background.

**Supplementary Table 4| Convergent mutations in the spike protein.** List of all mutations in the spike protein that converged at the nucleotide level between the evolved lines CoV-2-D and Cov-2-G.

**Supplementary Table 5| Convergent mutations in the N, M and E proteins.** List of all mutations in the N, M and E proteins, that converged at the nucleotide level between the evolved lines CoV-2-D and Cov-2-G.

**Supplementary Table 6| Mutations in common with the natural SARS-CoV-2.** List of non-synonymous mutations in Spike that emerged during our experiment in the CoV-2-D and Cov-2-G backgrounds or in the mutator lines and that were also observed in the natural population of SARS-CoV-2 (until the 24th of October 2021; https://nextstrain.org/ncov/gisaid/global). Among these, we observed the mutations H655Y (present in the variant of concern Gamma, lineage P.1, originated in Brazil), D215G (present in the variant of concern Beta, lineage B.1.351, firstly identified in South Africa) and D253G (found in lineage B.1.426, mostly detected in the US) (**Fig. 5b**).
